# A PDLP-NHL3 complex integrates plasmodesmal immune signaling cascades

**DOI:** 10.1101/2022.09.20.508228

**Authors:** Estee E. Tee, Matthew G. Johnston, Diana Papp, Christine Faulkner

## Abstract

The plant immune system relies on the perception of molecules that signal the presence of a microbe threat. This triggers signal transduction that mediates a range of cellular responses via a collection of molecular machinery including receptors, small molecules, and enzymes. One response to pathogen perception is the restriction of cell-to-cell communication by plasmodesmal closure. We previously found that while chitin and flg22 trigger specialized immune signaling cascades in the plasmodesmal plasma membrane, both execute plasmodesmal closure via callose synthesis at the plasmodesmata. Therefore, the signaling pathways ultimately converge at or upstream of callose synthesis. To establish the hierarchy of signaling at plasmodesmata and characterize points of convergence in microbe elicitor-triggered signaling, we profiled the dependence of plasmodesmal responses triggered by different elicitors on a range of plasmodesmal signaling machinery. We identified that, like chitin, flg22 signals via RBOHD to induce plasmodesmal closure. Further, we found that PDLP1, PDLP5 and CALS1 are common to microbe- and SA-triggered responses, identifying PDLPs as a candidate signaling nexus. To understand how PDLPs relay a signal to CALS1, we screened for PDLP5 interactors and found NHL3, which is also required for chitin-, flg22- and SA-triggered plasmodesmal responses and PDLP-mediated activation of callose synthesis. We conclude that a PDLP-NHL3 complex acts as an integrating node of plasmodesmal signaling cascades, transmitting multiple immune signals to activate CALS1 and plasmodesmata closure.

**Significance Statement:** Plants close plasmodesmata to restrict cell-to-cell communication after pathogen perception and in response to a range of stresses. All these stimuli trigger callose deposition at the plasmodesmal neck, suggesting a convergence of signaling. We have defined the hierarchy of molecular components and signals required to mediate plasmodesmal closure in immune responses, identifying a PDLP-NHL3 complex as a critical node that integrates multiple signaling cascades to regulate plasmodesmata.

## Introduction

The innate immune system of plants relies in part on the perception of invading pathogens at the cell surface. Cell-surface perception initiates signal transduction to induce cellular responses via molecular machinery that includes receptor kinase complexes, secondary messengers, and enzymes. This machinery can function across different subcellular structures, cells, and tissues, and be co-opted to mediate diverse defense responses. A single stimulus can intiate multiple signaling cascades: for example, receptor activation can trigger diverse responses that include a burst in production of reactive oxygen species (ROS) and changes to gene expression. Further, different stimuli can independently initiate cascades that converge on common responses such as the activation of callose synthesis in different cellular locations triggered by a range of microbial elicitors. By understanding the hierarchy and relationship of different signaling cascades, we can reveal the immune signaling network and identify critical nodes.

The microbe-associated molecular pattern (MAMP) elicitors chitin and flg22 induce plasmodesmata closure (1–3) via the activity of specific receptor complexes and modulators. Some of this machinery acts exclusively in plasmodesmata, establishing independence from signaling in the plasma membrane. For chitin-triggered plasmodesmal closure, specificity is conferred by the LYSM-CONTAINING GPI-ANCHORED PROTEIN 2 (LYM2) which acts independently of the canonical, plasma membrane located chitin receptor, CHITIN ELICITOR RECEPTOR KINASE 1 (1, 3). By contrast, flg22-triggered plasmodesmal closure depends on the canonical flagellin receptor FLAGELLIN SENSING 2, but is specifically mediated downstream by the plasmodesmata-localized calcium responsive protein, CALMODULIN-LIKE 41 (CML41) (1, 2). However, both chitin- and flg22-triggered plasmodesmata closure occurs via callose synthesis suggesting these specific cascades converge.

Plasmodesmal closure in response to a range of pathogen and stress elicitors is ultimately executed by callose deposition at the plasmodesmata neck, produced by callose synthases (4). PLASMODESMATA LOCATED PROTEIN 5 (PDLP5) positively regulates callose deposition via CALLOSE SYNTHASE 1 (CALS1) in response to the defense hormone salicylic acid (SA) (5), but the components that specifically mediate plasmodesmal callose deposition in response to chitin and flg22 are not yet known. Further, the observations that SA and ROS produced by the NADPH oxidase RESPIRATORY BURST OXIDASE HOMOLOGUE D (RBOHD) trigger plasmodesmal responses identifies that secondary messengers common to a range of immune responses also play specific roles in the activation of callose synthesis at plasmodesmata. Therefore, characterization of the signaling pathways triggered by different elicitors at plasmodesmata will define nodes that integrate distinct signals to enable a common response, and establish how specific responses are executed at plasmodesmata.

We sought to leverage our knowledge of plasmodesmal machinery that functions in specific responses to determine the relationships between signaling cascades triggered by different immune elicitors. We found that flg22, like chitin, elicits a cascade that signals via RBOHD, and that PDLP1, PDPL5 and CALS1 are common to microbe elicitor- and SA-triggered responses. To dissect the possibility that PDLPs act as a signaling nexus and understand how they activate callose synthesis, we identified NDR1 (nonrace-specific disease resistance)/HIN1 (hairpin-induced)- like protein 3 (NHL3) as a PDLP5 interactor. Specifically, NHL3 is a key plasmodesmal component required for PDLP-mediated activation of callose synthesis. The data support a model in which NHL3 interacts with PDLPs to form a central integrator of plasmodesmal immune signaling, transmitting information to activate CALS1 and plasmodesmata closure.

## Results

### Chitin- and flg22-triggered plasmodesmata closure signal via RBOHD to a downstream nexus requiring PDLP5

To investigate whether and where plasmodesmal signaling pathways converge upstream of callose deposition, we surveyed known plasmodesmal signaling components for their role in response to specific elicitors. Focusing initially on chitin- and flg22-triggered plasmodesmal responses, we hypothesized that as chitin-triggered plasmodesmal closure is mediated by activation of RBOHD (3), and that flg22 triggers RBOHD activation (6), RBOHD might also mediate flg22-triggered plasmodesmal signaling. Thus, we assayed for flg22-triggered plasmodesmal closure in *rbohD* mutants using microprojectile bombardment assays and found that *rbohD* cannot close plasmodesmata in response to flg22 (Fig. 1a), demonstrating that flg22-triggered plasmodesmal closure signals via RBOHD. For chitin, RBOHD dependence is linked to phosphorylation of S133 and S347 (3) and we therefore assayed for flg22-triggered plasmodesmal closure in *rbohD* mutants complemented with alleles of RBOHD in which active phosphorylation sites (6) are mutated: RBOHD_S163A_, RBOHD_S133A_, RBOHD_S343A/S347A_ or RBOHD_S39A/S339A/S343A_. This revealed that neither RBOHD_S133A_ nor RBOHD_S343A/S347A_ complemented the loss of *rbohD* dependent flg22 induced plasmodesmal closure (Fig. S1) and that S133 and S347 of RBOHD are required for both flg22- and chitin-triggered plasmodesmal response.

**Fig. 1.**
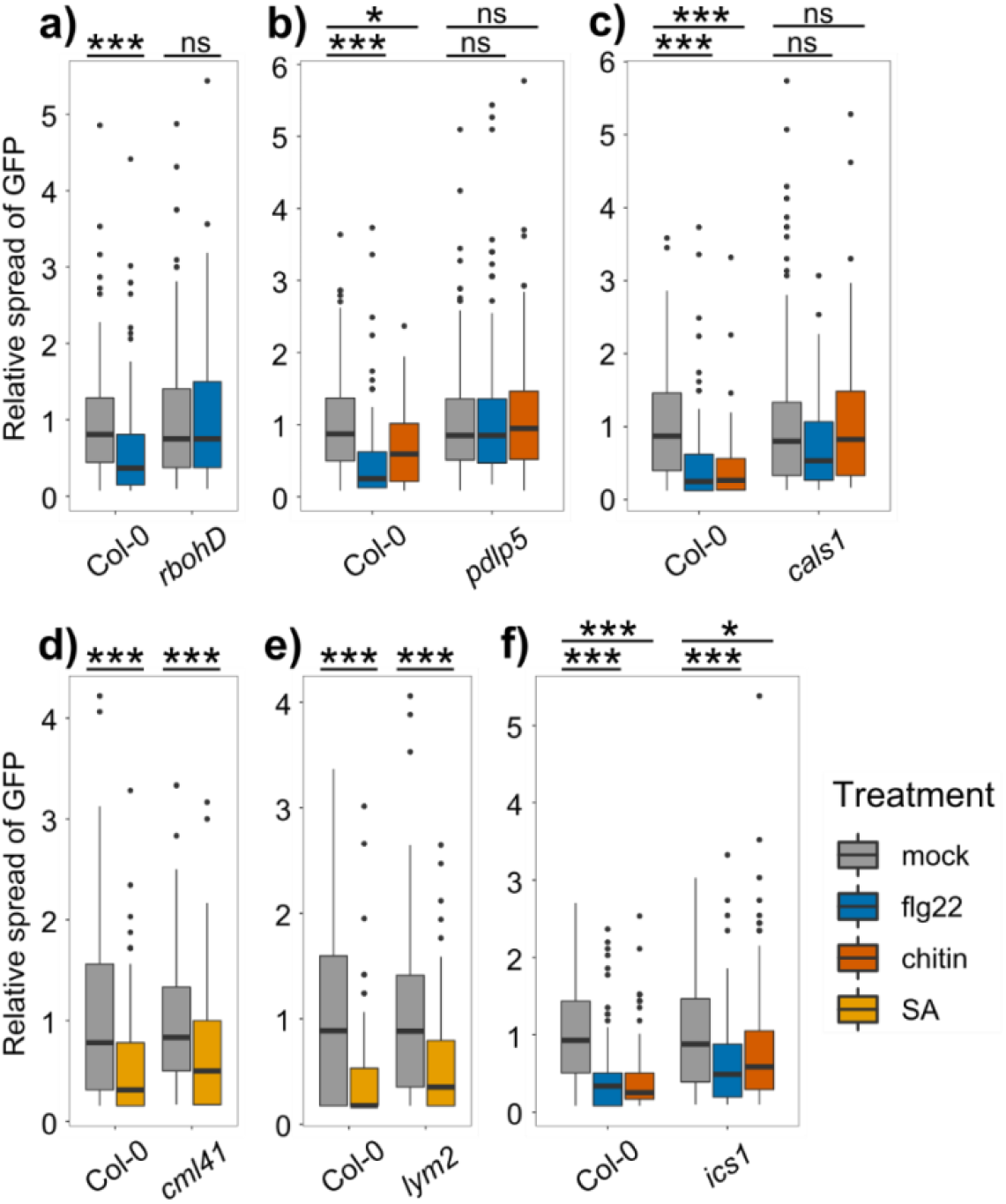
Hierarchical dissection of plasmodesmata immune responses reveals common signaling components. Relative GFP movement into neighboring cells following microprojectile bombardment. a) flg22 reduces GFP movement to neighboring cells in Col-0 but not in *rbohD* (n≥103 bombardment sites). b) chitin and flg22 reduce GFP movement to neighboring cells in Col-0 but not in *pdlp5* (n≥58 bombardment sites). c) chitin and flg22 reduce GFP movement to neighboring cells in Col-0 but not in *cals1* (n≥120 bombardment sites). SA reduces the movement of GFP to neighboring cells in Col-0, *cml41* (d, n≥119 bombardment sites) and *lym2* (e, n≥117 bombardment sites). f) chitin and flg22 reduce GFP movement to neighboring cells in Col-0 and *ics1* (n≥114 bombardment sites). Asterisks indicate statistical significance compared to the mock treatment within a genotype: \**p*<0.05, \*\*\**p*<0.001.

SA-triggered plasmodesmata closure is independent of RBOHD (5) but, like chitin and flg22, induces callose deposition. Chitin- and flg22-induced plasmodesmal closure occurs via callose deposition within 30 minutes (2, 3) and so to determine if SA might plausibly exploit a similar signaling mechanism, we assayed for SA-induced callose deposition at plasmodesmata. Quantifying aniline blue stained plasmodesmata associated callose, we observed that SA induces plasmodesmal callose deposition within 30 minutes of treatment (Fig. S2). Therefore, SA responses have a similar temporal profile to chitin and flg22 responses, allowing for the possibility that SA, chitin and flg22 induce plasmodesmal closure via a similar mechanism.

To explore whether flg22, chitin and SA signaling converge on the same pathway upstream of callose deposition, we tested the requirement for known mediators of SA-triggered plasmodesmata closure, PDLP5 and CALS1, in chitin and flg22 responses. Using microprojectile bombardment assays, we found that neither *pdlp5* (Fig. 1b) nor *cals1* (Fig. 1c) mutants closed their plasmodesmata in response to flg22 or chitin. As CALS1 produces callose and must lie at the endpoint of the signaling pathways, we conclude that flg22, chitin, and SA signaling at plasmodesmata converge downstream of RBOHD, and at, or upstream of, PDLP5.

### Immune-associated salicylic acid production is not required for elicitor-induced plasmodesmal closure

As SA is known to be synthesized in response to flg22 treatment (7, 8), we aimed to determine whether SA mediates flg22- and chitin-triggered plasmodesmal closure. Given that *rbohD* exhibits normal plasmodesmal closure response to SA (5), and that *RBOHD* is required for both chitin- and flg22, we reasoned that a role for SA-triggered and/or mediated responses would lie downstream of LYM2 and CML41. To test this, we first generated a null *cml41* mutant (Fig. S3) by CRISPR/Cas9 gene editing that is perturbed in flg22-induced plasmodesmal closure (Fig. S3d) and found that this mutant can close its plasmodesmata in response to SA (Fig. 1d). Likewise, *lym2* (Fig. 1e) shows wild-type-like SA-induced plasmodesmata closure.

To assay for a direct role of SA in chitin- and flg22-triggered plasmodesmal closure, we asked whether these elicitors trigger plasmodesmal closure if SA synthesis is perturbed. Thus, we assayed for plasmodesmal responses in the ISOCHORISMATE SYNTHASE 1 (*ics1*) mutant, in which pathogen-induced SA synthesis and accumulation is significantly reduced. We observed that *ics1* plasmodesmata can close in response to chitin and flg22, and therefore chitin- or flg22-triggered SA production is not necessary for plasmodesmal signaling (Fig. 1f). Collectively, these data suggest SA itself does not act as a secondary messenger within microbe-triggered plasmodesmal cascades, but rather initiates a signaling pathway that converges with other responses.

### PDLP1 is required for immune elicitor-triggered plasmodesmata closure

PDLP1 positively regulates callose deposition at plasmodesmata and at the downy mildew haustorial membrane (9) but has not been associated with plasmodesmal responses. However, it has been shown that PDLP5 and PDLP1 can interact (10), suggesting PDLP1 might co-operatively function in the same responses as PDLP5. To test this, we assayed the *pdlp1* mutant for its ability to respond to flg22 and SA and found neither induced plasmodesmal closure (Fig. S4). Therefore, PDLP1 is also required for elicitor-triggered signaling cascades in plasmodesmata, supporting a model in which PDLPs are common to multiple signaling pathways and function downstream of, or at, the point at which these signals converge.

### The NDR1/HIN1-like protein NHL3 interacts with PDLP5 at plasmodesmata

Having established that PDLP5 functions in MAMP- and SA-triggered plasmodesmal signaling, and considering it is a positive regulator of callose deposition (5, 11), we hypothesized that PDLP5 is a component of a central node in immune-triggered plasmodesmal signaling cascades. To characterize its function in this context, we aimed to identify PDLP5 interactors that might link PDLPs to callose synthase activity. Thus, we commissioned a split-ubiquitin screen of an *Arabidopsis thaliana* (*Arabidopsis*) cDNA library with a PDLP5 bait (DualSystems Biotech, Switzerland; now Hybrigenics Services, France). The screen identified 65 prey hits and we refined this list based on their presence in plasmodesmal proteomes of *Arabidopsis* suspension culture cells (12, 13), producing 13 prey proteins (Table S1). NHL3 (AT5G06320) is present in both proteomes and is a membrane-associated protein like PDLP5. Thus, we chose NHL3 for further characterization as a likely PDLP5 interactor.

We first examined the subcellular localization of NHL3 to determine whether a *bona fide* interaction with PDLP5 is plausible. We C-terminally tagged NHL3 with mCherry and observed that NHL3-mCherry is located in the plasma membrane, as previously reported ((14, 15), Fig. 2a). However, NHL3-mCherry fluorescence also accumulated in puncta along the plasma membrane that co-localized with both PDLP5-eGFP and PDLP1-eGFP, indicating that it accumulates in plasmodesmata (Fig. 2a).

**Fig. 2.**
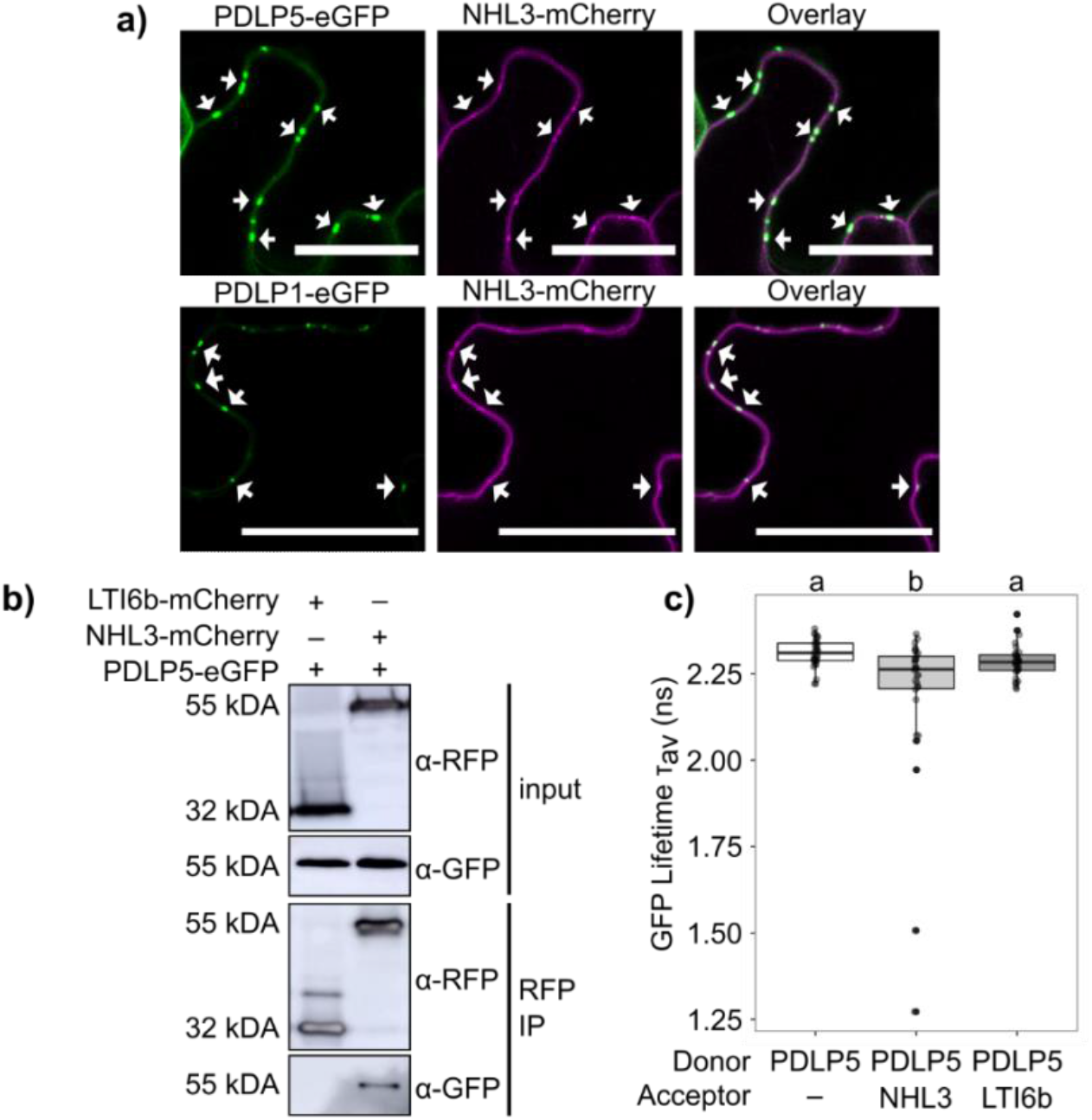
NHL3 localizes to plasmodesmata and interacts with PDLP5. a) Confocal micrographs of *N. benthamiana* transiently expressing PDLP5-eGFP (green) or PDLP1-eGFP (green) and NHL3-mCherry (magenta). Arrows point to example plasmodesmata where fluorescence overlays. Scale bars are 25 μm. b) Western blot analysis of immunoprecipitated (IP) protein extracts from *N. benthamiana* expressing PDLP5-eGFP with LTI6b-mCherry or NHL3-mCherry; representative of three independent repeats. c) FRET-FLIM analysis of transiently expressed constructs in *N. benthamiana* of PDLP5-eGFP alone (PDLP5), and in the presence of acceptors NHL3-mCherry (NHL3) or LTI6b-mCherry (LTI6b). Independent factor ‘Genes Expressed’ was determined as significant (ANOVA; F = 6.7, df = 2, *p*<0.01), with significant differences denoted by a and b, *p*<0.05. Each sample is represented by six measurements from six biological replicates.

To confirm that NHL3 interacts with PDLP5, we first used targeted co-immunoprecipitation (co-IP). NHL3-mCherry, or the negative control LTI6b (LOW TEMPERATURE INDUCED PROTEIN 6b)-mCherry, was transiently co-expressed with PDLP5-eGFP in *N. benthamiana* leaves. Immunoprecipitation of mCherry-tagged proteins showed that PDLP5-eGFP co-immunoprecipitated with NHL3-mCherry, but not with LTI6b-mCherry (Fig. 2b), supporting the hypothesis that PDLP5 specifically associates with NHL3.

To further test this association, we used Förster resonance energy transfer–fluorescence lifetime imaging (FRET-FLIM) analysis. As NHL3 has been differentially predicted to carry one or two transmembrane domains (14), and a strategy for epitope tagging to allow FRET was unclear, we first resolved the membrane topology of NHL3 with bimolecular fluorescence (BiFC) complementation by transient expression in *N. benthamiana*. NHL3 was either N- or C-terminally tagged with the N-terminus of yellow fluorescent protein (YFP^N^) and co-expressed with a cytosolic (YFP^C^) (Fig. S5a, b) or a secreted (SP-YFP^C^) C-terminus of YFP (Fig. S5c, d). YFP fluorescence was visible in the positive control combinations: cytosolic YFP^C^ co-expressed with cytosolic YFP^N^ (Fig. S5e), and SP-YFP^C^ with SP-YFP^N^, (Fig. S5f). YFP fluorescence was similarly detected when NHL3-YFP^N^ or YFP^N^-NHL3 was co-expressed with cytosolic YFP^C^ (Fig. S5a and S5b), but not with SP-YFP^C^ (Fig. S5c and S5d). Therefore, both termini of NHL3 are cytosolic facing.

We assayed for interaction between NHL3-mCherry and PDLP5-eGFP fusion proteins using FRET-FLIM. As a negative control, we fused LTI6b to mCherry and an *ACT2* promoter to drive expression (*pACT2::LTI6b-mCherry*) and found that, unlike previous reports (16), LTI6b localizes to the plasma membrane and was visible in puncta in the plasma membrane coincident with PDLP5-eGFP (Fig. S6). This allowed us to resolve our FRET-FLIM analysis to plasmodesmata within the plasma membrane. We found that the fluorescence lifetime of PDLP5-eGFP was reduced at plasmodesmata in the presence of NHL3-mCherry, but not LTI6b-mCherry (Fig. 2c), indicating that FRET occurs between PDLP5 and NHL3. Alongside co-IP data and the initial split ubiquitin screen, this supports a model in which PDLP5 and NHL3 form a complex for plasmodesmal signaling.

### NHL3 is required for immune-associated plasmodesmata closure

The *NHL* gene family has homology to *NON-RACE SPECIFIC DISEASE RESISTANCE* (*NDR1*) and the *TOBACCO HAIRPIN-INDUCED* (*HIN1*) genes (17). However, while *HIN1* and *NHL* members have been associated with various aspects of plant defense and stress responses (14, 18–21), the specific molecular functions of NHLs are unknown. Given the plasmodesmal localization of NHL3, and its interaction with PDLP5, we investigated the plasmodesmal responses of *nhl3* mutants to chitin, flg22, H_2_O_2_ and SA. Plants carrying the *nhl3* mutant alleles, *nhl3*-1 and *nhl3*-2, could not close their plasmodesmata in response to chitin or flg22 (Fig. 3a). Furthermore, while *nhl3* mutants were able to close their plasmodesmata in response to H_2_O_2_, they could not in response to SA (Fig. 3b), phenocopying the *pdlp5* and *cals1* mutants (5, 22). Thus, NHL3 is required for elicitor-triggered plasmodesmal responses.

**Fig. 3.**
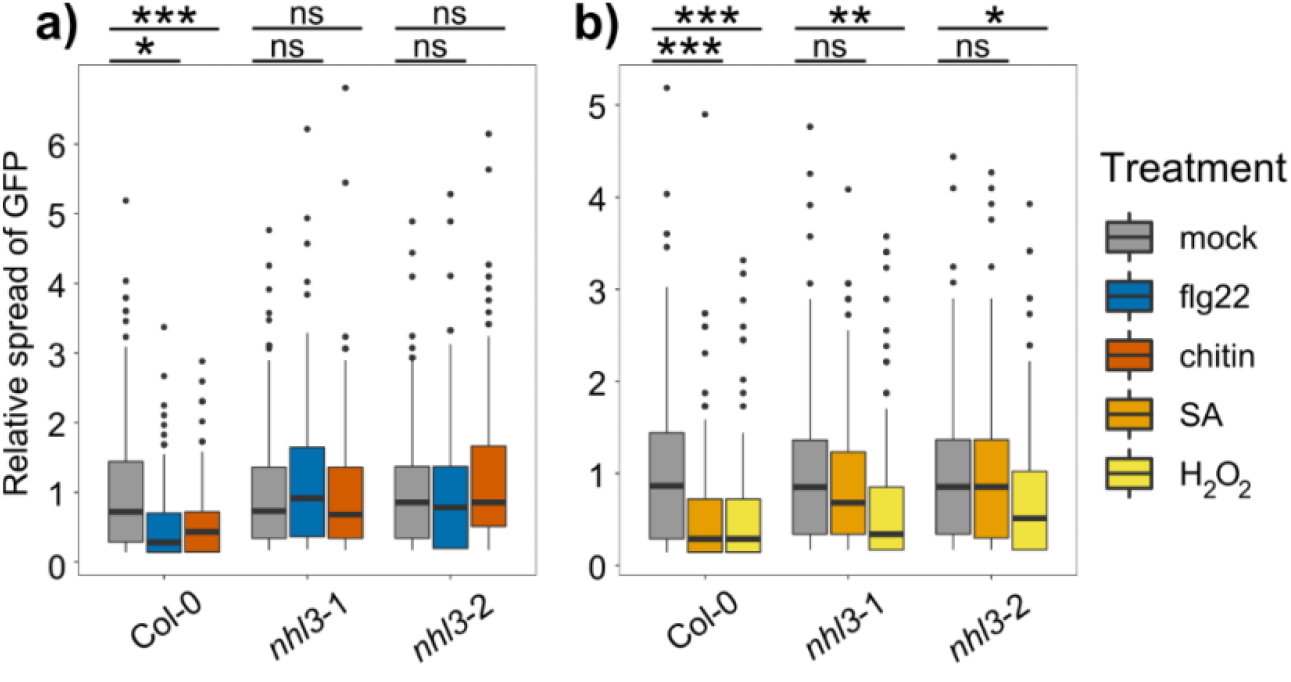
NHL3 functions in plasmodesmal immune signaling. Relative GFP movement into neighboring cells following microprojectile bombardment. a) flg22 and chitin reduces GFP movement to neighboring cells in Col-0, but not in *nhl3*-1 and *nhl3*-2. b) H_2_O_2_ reduces GFP movement to neighboring cells in Col-0, *nhl3*-1 and *nhl3*-2. SA reduces GFP movement to neighboring cells in Col-0 but not in *nhl3*-1 and *nhl3*-2 (n≥ 118 bombardment sites). Asterisks indicate statistical significance compared to the mock treatment within a genotype: \**p*<0.05, \*\**p*<0.01, \*\*\**p*<0.001.

### NHL3 positively regulates callose deposition at plasmodesmata

Reasoning that NHL3 likely co-operates with PDLP5 to positively regulate callose deposition at plasmodesmata, we assayed for the impact of NHL3 over-expression on callose deposition at plasmodesmata using quantitative imaging of aniline blue stained callose. When *NHL3* and *PDLP5* were transiently expressed individually or together in *N. benthamiana*, there was a significant increase in total aniline blue fluorescence in a plasmodesmal callose deposit compared to the infiltration control (Fig. 4a), independent of whether NHL3 was N- or C-terminally tagged (Fig. S7). When *NHL3* and *PDLP5* were co-expressed, we did not detect a further increase in callose per plasmodesmal deposit, but we did detect an increase in the number of deposits detected per field of view compared to when both proteins were individually expressed (Fig. S8). This might arise from more plasmodesmata producing callose above the detection threshold and indicate increased callose synthesis. These data support a function of NHL3 in co-operating with PDLP5 to increase the activity of callose synthases, thereby positively regulating callose deposition at plasmodesmata.

**Fig. 4.**
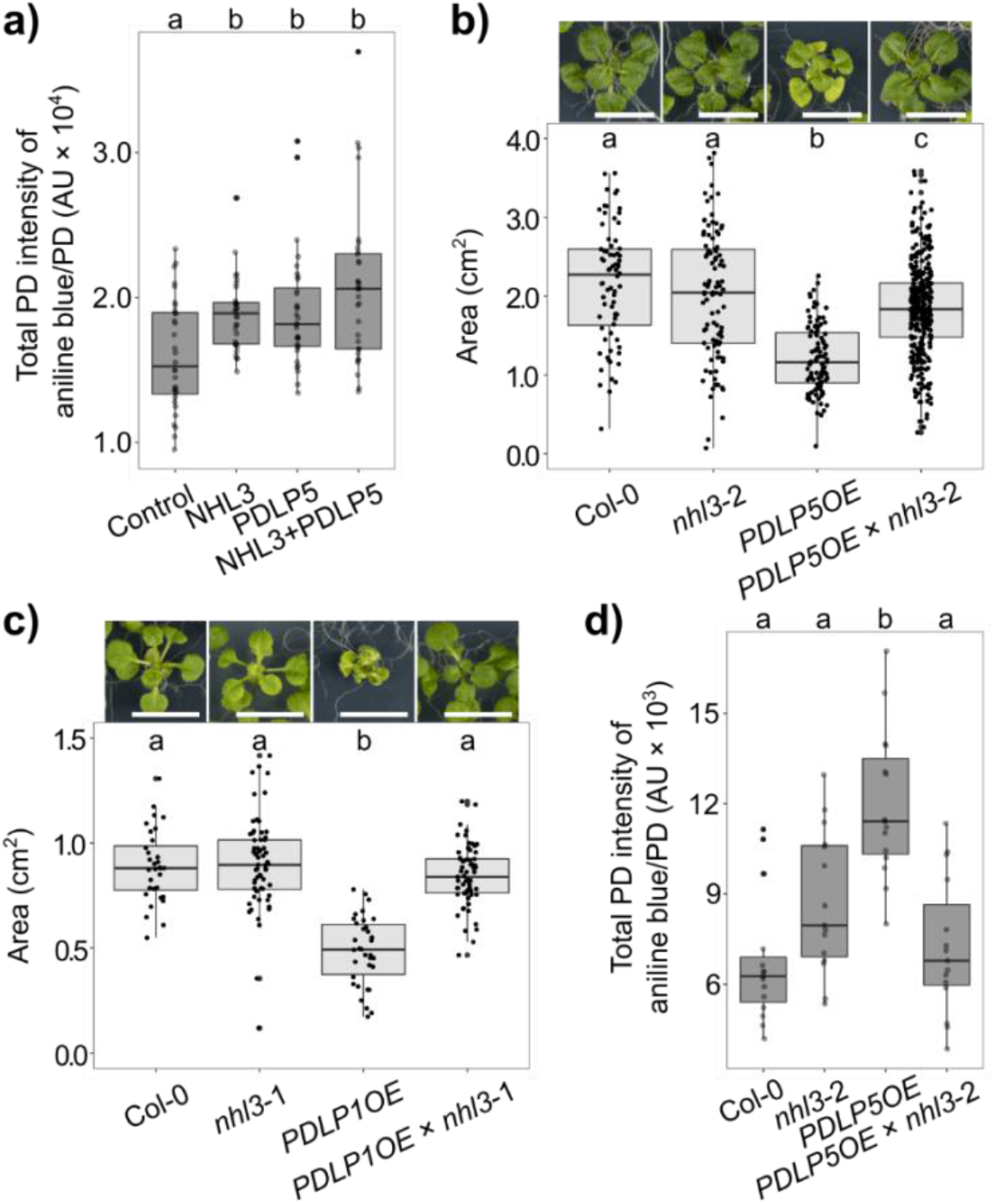
NHL3 positively regulates callose deposition at plasmodesmata and is essential for PDLP function. a) Quantification of aniline blue-stained plasmodesmata-associated callose using automated image analysis in *N. benthamiana* transiently expressing p19 only (Control), NHL3-mCherry (NHL3), PDLP5-eGFP (PDLP5) or NHL3-mCherry and PDLP5-eGFP (NHL3+PDLP5). Combined data from two independent experiments, with four images from four plants taken per construct combination in each experiment. Independent factor ‘Gene Expressed’ was determined as significant (ANOVA: F = 8.6, df = 3, *p*<0.0001), with significant differences denoted by a and b, *p*<0.05. b) Whole rosette area of 21-day-old plants of Col-0, *nhl3*-2, *PDLP5OE* and *PDLP5OE* × *nhl3*-2. Combined data from two independent experiments with random effects being ‘experiment’ and ‘the different individual plates plants were grown’. Independent factor ‘Genotype’ was determined as significant (ANOVA: F = 80.3, df = 3, *p*<0.0001), with significant differences denoted by a, b and c, *p*<0.0001. For Col-0, *nhl3*-2, *PDLP5OE* and *PDLP5OE* × *nhl3*-2, n = 72, 97, 100 and 318, respectively. Scale bar is 1 cm. c) Whole rosette area of 21-day-old Col-0, *nhl3-*1, *PDLP1OE* and *PDLP1OE* × *nhl3-*1. Independent factor ‘Genotype’ was significant (ANOVA: F = 45.4, df = 3, *p*<0.0001), with significant differences denoted by a and b, *p*<0.0001. N for Col-0, *nhl3*-1, *PDLP1OE* and *PDLP1OE* × *nhl3*-1 is 32, 64, 34 and 63, respectively. Scale bar is 1 cm. d) Quantification of aniline blue stained plasmodesmata-associated callose using automated image analysis in 3-week-old *Arabidopsis* leaves, with three images for five plants taken per genotype. Independent factor ‘Genotype’ was determined as significant (ANOVA: F = 14.0, df = 3, *p*<0.0001), with significant differences denoted by a and b, *p*<0.01.

### PDLP-activated callose synthesis requires NHL3

To dissect the hierarchy of the relationship between PDLP5 and NHL3 we asked whether PDLPs require NHL3 to function. For this, we used plants that overexpress *PDLP5* (*PDLP5OE*, (11)) and *PDLP1-GFP* (*PDLP1OE* (4)), which are both phenotypically dwarfed in comparison to wild-type plants. We crossed *PDLP5OE* with *nhl3*-2 (*PDLP5OE* × *nhl3*-2) and *PDLP1OE* with *nhl3-1* (*PDLP1OE* × *nhl3-1*) and found that, in both cases, mutations of *nhl3* reverted the dwarf phenotypes of *PDLP* over-expressors (Fig. 4b, Fig. 4c, Fig. S9). In light of this dwarf reversion, and considering PDLP1, PDLP5 and NHL3 are associated with positive regulation of callose at plasmodesmata ((4, 9, 11), Fig. 4a), we assayed for callose deposition at plasmodesmata in *PDLP5OE* × *nhl3*-2 lines, relative to *PDLP5OE*. We confirmed that *PDLP5OE* plants exhibit high callose deposition at plasmodesmata in our growth conditions and found that this was reverted to wild-type levels in *PDLP5OE* × *nhl3*-2 plants (Fig. 4d). Overall, this suggests that NHL3 is common to downstream functions of both PDLP1 and PDLP5, integrating a range of signals to induce plasmodesmal callose deposition.

## Discussion

Plants deploy a suite of signaling cascades with specific molecular components to initiate and execute defense responses against invading pathogens. These cascades, and the machinery they require, can be specific to subcellular structures such as is observed in plasmodesmata, where microbe elicitors can trigger discrete responses. Despite the specificity in signaling observed in plasmodesmata, pathways initiated by different elicitors converge on the common response of callose synthesis to mediate plasmodesmal closure. Thus, plasmodesmal responses likely converge at a node that integrates multiple independent inputs.

To identify and characterize this node, we systematically determined consecutive steps in the signaling pathways for elicitor-induced plasmodesmal closure. This allowed us to determine that immune-associated signaling cascades progressively converge on a single node involving PDLPs and NHL3, upstream of CALS1 activation (Fig. S10). Our data indicate that at the top of the signaling cascade, chitin and flg22 both relay signals via specific receptors and molecules to RBOHD ((3), Fig. 1a, S1). As an NADPH oxidase, RBOHD produces ROS that act as small molecule transmitters of plasmodesmal responses. How ROS are perceived and transmit information within this cascade remains an open question and the observation that hydrogen peroxide can induce plasmodesmata closure via an independent pathway (5) complicates experimental dissection of this signaling step without a candidate receiver. However, increasing knowledge of proteins present in plasmodesmata and the role which they play will likely reveal candidate components to fill this gap.

Our data suggest that SA elicits plasmodesmal responses by a signaling cascade that converges on microbial-triggered responses downstream of RBOHD. Our observation that SA-induced plasmodesmal closure occurs within 30 minutes (Fig. S2), as well as its dependence on *PDLP1* (Fig. S4) which is not transcriptionally induced by SA (23), challenges the hypothesis that SA-triggered plasmodesmal closure occurs only via SA-induced expression of *PDLP5*. However, how SA might elicit plasmodesmal closure via another mechanism is not currently clear.

Leveraging our new understanding of PDLPs as a common component of different immune-associated plasmodesmal responses, we identified NHL3 as a co-operative interactor of PDLP5. We found that NHL3 positively regulates PDLP5- and PDLP1-mediated callose deposition, and functions in elicitor-triggered plasmodesmal responses. Given that PDLP1 and PDLP5 interact (10) we propose they function in complex with NHL3, forming an integrator of immune signal inputs. PDLPs are only found in seed plants (24–28), indicating a recent emergence in comparison to NHL related proteins which have orthologues to proteins found in plasmodesmal fractions of the earlier land plant *Physcomitrium patens* (26, 29); this allows speculation that PDLPs were co-opted into a pre-existing complex of NHLs with callose synthases to enable an extra regulatory mechanism for plasmodesmata responses.

How NHL3 mediates signaling is not clear as its only annotated domain is a LATE EMBRYOGENESIS ABUNDANT (LEA) protein LEA_2 domain found close to the C-terminus, which has no known function. LEA_2 domain proteins are produced during responses to dehydration or hyperosmotic stress, and have been postulated to stabilize macromolecules against freezing, dehydration, ionic or osmotic stress (30). They have a hydrophobic (31), glycine-rich chemical composition (32), and are generally predicted to be intrinsically unstructured proteins with a left-handed polyproline II helix confirmation (31, 33–35). This disordered confirmation is thought to form tight hydrogen-bond networks in concert with carbohydrates, providing stability to macromolecular structures (30, 36). Thus, any role for NHL3 in plasmodesmal signaling might relate to recruitment, aggregation or stabilization of signaling or response components. Given that the NHL protein family has many members in Arabidopsis, and that another member has also been found to have plasmodesmata-associated localization and functions (37), the mode of action of NHLs might regulate plasmodesmata in diverse response contexts.

PDLPs have two extracellular DOMAIN OF UNKNOWN FUNCTION (DUF) 26 domains. Structural analysis of the PDLP5 DUF26 domains suggests these are lectin dimers, which could form an extended binding cleft for a carbohydrate polymer (24). While the PDLP5 DUF26 domain did not bind to a range of cell wall-derived carbohydrates (24), callose binding has not been tested. Considering the LEA_2 domain sits on the cytosolic C-terminus of NHL3, it seems possible that NHL3 might serve as a scaffold for PDLP5 and CALS1, thereby stabilizing the machinery required for callose deposition in plasmodesmata. Indeed, such a model might challenge the notion of a linear signaling pathway, in that PDLP1/5 and NHL3 might not only activate callose synthesis but could also bind and stabilize callose at the site of synthesis via the DUF26 domains of PDLPs. The observation that *PDLP1* and *PDLP5* overexpression phenotypes are reverted to wild-type levels in the absence of *NHL3* (Fig. 4b-d) suggests NHL3 either acts downstream of PDLPs or that they are co-dependent. A phloem-specific PDLP-CALS interaction has recently been identified (38); however, resolving whether PDLP5 or NHL3 directly activates CALS1, or whether they transmit a signal to an intermediary component, will further dissect how the PDLP5/NHL3 complex acts to regulate callose synthases.

How plasmodesmata specifically respond to a range of immunity- and stress-associated elicitors is a critical question that underpins how organism-level responses are regulated in a multicellular context. Further, how immune signals are localized, specialized, and integrated is key to mapping the molecular signaling networks that orchestrate cellular responses to pathogens. Characterization of plasmodesmata signaling has revealed that a PDLP-NHL3 complex integrates different signaling cascades to activate callose synthesis and close plasmodesmata during immune responses. Thus, we have uncovered a mechanism by which diverse signals are integrated to produce a single response, further characterizing the signaling cascades that regulate cell-to-cell connectivity during immune responses. How NHL3 functions, and the full range of signals it can integrate, in addition to resolving how small molecules such as ROS and SA are received in plasmodesmal responses, remain questions to be addressed to understand the molecular dynamics of plasmodesmal closure and callose synthesis across plant responses.

## Materials and Methods

### Extended Methods and Materials are available in *SI Appendix*

#### Plant material

*Arabidopsis thaliana* wild-type used in this study is Columbia-0 (Col-0). Mutants and transgenic lines are in the Col-0 background. Lines used in this study are *lym2* (*lym2*-1, SAIL_343_B03, (1)); *ics1* (*sid2*-1, (39)); *pdlp5* (*pdlp5*-1, SAIL_46_E06, (11)); *cals1* (*cals1*-1, SALK_142792, (40)); *rbohD* (*atrbohD*, (41)); RBOHD_S133A_ (3); RBOHD_S163A_ (3); RBOHD_S343A/S347A_ (6); RBOHD_S39A/S339A/S343A_ (6)); *nhl3*-1 (SALK_035427, (42)); *nhl3*-2 (SALK_150318, (43)); PDLP5OE (*35S:PDLP5*, (11)); PDLP1OE (*35S::PDLP1-GFP*, (4)). The generation of the CRISPR/Cas9 *cml41* null mutant is outlined in Extended Material and Methods (*SI Appendix*) and Fig. S3. For bombardment assays and callose staining, *Arabidopsis* plants were grown on soil with 10 h light at 22 °C. *Nicotiana benthamiana* plants were grown on soil with 16 h light at 23 °C, with leaves of 4-week-old plants used for transient expression.

#### Microprojectile bombardment assays

Bombardment assays were performed as described in (44). Bombardment sites were imaged 16 hours after bombardment using a confocal microscope (LSM Zeiss 800) with a 20× water dipping objective (W N-Achroplan 20×/0.5).

#### Plasmodesmata callose staining and quantification

The 7^th^ or 8^th^ *Arabidopsis* leaf or *N. benthamiana* leaf was infiltrated with 1% or 0.1% aniline blue in PBS buffer (pH 7.4) respectively. Plasmodesmata callose deposits were imaged from the abaxial side by a confocal microscope (LSM Zeiss 800) using a 63× water immersion objective (Plan-Apochromat 63×/1.0 M27). Aniline blue was excited with a 405 nm UV laser and collected at 430-550 nm. Z-stacks from multiple areas (specified in figure legends) per leaf were collected. We used an automated image analysis pipeline to quantify aniline blue stained plasmodesmata, available at (45).

#### Split-ubiquitin screen

The split-ubiquitin screen was performed by DualSystems Biotech, Switzerland (now known as Hybrigenics Services, France).

#### Co-immunoprecipitation

Protein extraction and co-IP was performed as described (3) with slight modifications (SI Appendix). Immunoblotting was carried out with anti-GFP (Roche, 11814460001; 1:1000) or anti-RFP-biotin (Abcam, ab34771, 1:2000) primary antibodies, and anti-mouse IgG (Sigma, A0168; 1:10000) or anti-rabbit-HRP (Sigma, A0545, 1:10000) secondary antibodies.

#### Confocal microscopy

Transiently expressed constructs for co-localization and split-YFP analysis were imaged by confocal microscopy (LSM Zeiss 800) two days post-infiltration using a water immersion 63× objective (C-Apocromat 63×/1.20WKorr UV VIS IR-water). GFP/YFP was excited with a 488 nm argon laser and collected at 505-530 nm while mCherry was excited with a 561 nm DPSS laser and collected at 600-640 nm.

#### FRET-FLIM

Leaves of *N. benthamiana* transiently expressing constructs of interest were imaged using a STELLARIS 8 FALCON microscope (Leica Microsystems). The abaxial side of the leaf sample was imaged using a 63× objective (Leica, HC PL APO CS2 5 63×/1.20 Water). GFP was excited with a 488 nm WLL laser and collected at 503-530 nm. Acquisition was set at a mode of xyz, format size of 512 × 512, scan speed of 40 Hz, single line repetitions and a pixel dwell time of 3.16 μs. To reduce the risk of bleaching the sample, laser power was adjusted to ensure no more than 1.0 photon per pulse occurred. Data sets were acquired by scanning each image until a suitable number of photon counts per pixel were reached in the brightest channel, up to a maximum of 1000 counts per pixel. Data were analyzed using the LAS X software. Plasmodesmal regions of interest (ROI) were specified using the intensity threshold and manual selection. Using a two-exponential decay for GFP, a fit was generated and the mean τ amplitude-weighted GFP lifetime (in ns) for all plasmodesmata per image was calculated.

#### Rosette area phenotyping

For rosette area phenotyping and subsequent callose staining, *Arabidopsis* seeds were surface sterilized with 70% ethanol and 2% bleach with 0.02% Triton X-100, then washed with sterile dH_2_O and sown onto Murashige and Skoog medium 1% sucrose 0.8% agar plates 16 h light at 22 °C. Photos were taken 21 days post-germination for determination of rosette area. Individual plants were traced by hand and area of the rosette was measured in FIJI.

#### Statistical analyses

Data presented as boxplots have the middle line marking the median, the box indicating the upper and lower quartiles, and the whiskers showing the minimum and maximum values within 1.5× interquartile range. Data were analyzed using RStudio 2021.09.01 Build 351/R version 4.0.3.

## Supporting information

Combined Supplemental Information

## Acknowledgments

We acknowledge access to the John Innes Centre Bioimaging Facility and thank Dr Sergio Lopez for their assistance and training with the microscopes, as well as the John Innes Centre Scientific Photography Department. The *ics1* seeds were kindly provided by Jonathan Jones (The Sainsbury Laboratory) and the LTI6b cloning module was made by Sebastian Samwald (JIC). This work was funded by the Biotechnology and Biological Research Council (BBS/E/J/000PR9796), the European Research Council (grant 725459 ‘INTERCELLAR’) and a John Innes Foundation PhD studentship to MGJ.

## Author Contributions

E.E.T., M.G.J., D.P. and C.F. designed research; E.E.T., M.G.J., and D.P. performed research; E.E.T., M.G.J., D.P. and C.F. analyzed data; E.E.T. and C.F. wrote the paper.

## Competing Interest Statement

No competing interests to declare.

## Notes

### Competing Interest Statement

The authors have declared no competing interest.

