## Supplementary material for "A PDLP-NHL3 complex integrates plasmodesmal immune signaling cascades": Combined Supplemental Information

### Extended Material and Methods

*Microprojectile bombardment assays*

Leaves of 5- to 6-week-old *Arabidopsis* were used for assaying plasmodesmal permeability, with a minimum of four leaves used per genotype and treatment. Bombardment assays were performed using 1 nm gold particles (BioRad) coated with pB7WG2.0-GFP and pB7WG2.0-$RFP_{ER}$ using a Biolistic PDS-1000/HE particle delivery system (BioRad). Treatments [mock ($dH_2O$), chitin (500 µg/mL), flg22 (100 nM), SA (100 µM), or $H_2O_2$ (10 mM)] were syringe infiltrated two hours post-bombardment. To visualize bombardments, GFP was excited with a 488 nm argon laser and collected at 505-530 nm while mRFP was excited with a 561 nm DPSS laser and collected at 600-640 nm. The number of cells showing GFP was normalized to the mean of the mock-treated data within a genotype, and N stated in each legend is bombardment sites collected per genotype/treatment.

*Generation of CRISPR-Cas9* cml41 *null mutant*

A construct for CRISPR-Cas9 gene editing was assembled as described (1). sgRNAs targeting *CML41* were designed and assembled by PCR and Golden Gate cloning with a *Ubi10 promoter::Cas9::Nos* terminator cassette and a bialaphos resistance gene (*bar*) in a binary vector for plant expression (Fig. S3a). The target sgRNA binding site in the *CML41* gene resulted in a 193 bp deletion (Fig. S3a) and introduced a premature stop codon. The *cml41* deletion was confirmed in candidate knockouts by genotyping (Fig. S3b; using primers GACATCTCCATCCATGAGAAACC and TGGCAAATGTTCAGCAGAGC), and *CML41* transcript levels were assessed by RT-PCR. 10-day old seedlings were grown on Murashige and Skoog 1% sucrose 0.8% agar plates under 16 h light at 22 °C. Total RNA was isolated using an RNeasy® Plant Mini Kit (Qiagen), according to the manufacturer's instructions. RNA was treated with a DNase treatment using the 'Rigorous DNase treatment' protocol from the TURBO DNA-*free*$^{TM}$ Kit (Invitrogen). cDNA was synthesized from 1 µg of RNA using the High-Capacity cDNA Reverse Transcription Kit with RNase Inhibitor (Thermo Fisher Scientific). PCR amplification using a 1:10 dilution template from the synthesized cDNA, was performed with GoTaq Green Master Mix (Promega) using either 25 amplification cycles for the housekeeper gene *GAPC2* as a control (using primers TCGGAAGAATCGGTCGTTTGG and TGTATGTCATGTACTCGGTGG) or 35 amplification cycles for *CML41* (using primers TGATGACAAGAAGAGATCTTTACGG and CATCTCAAACGCTGTCTTCAA). RT-PCR analysis confirmed *cml41* mutants did not produce full length *CML41* transcript (Fig. S3c). Loss of flg22-induced plasmodesmata closure confirmed the *cml41* mutant phenotype (Fig. S3d).

*Cloning of plant expression constructs*

For transient expression of genes in this study (*PDLP5*, *NHL3*, *LTI6b*, *PDLP1*) coding sequences were cloned with internal BpiI/BsaI/Esp3I sites removed for Golden Gate assembly. All genes were fused with an epitope tag (eGFP, mCherry, $\text{YFP}^{N}$, $\text{YFP}^{C}$) and assembled for *in planta* expression with a 35S CaMV promoter (*p35S*) and a CaMV 35S or *heat shock protein 18.2* (*HSP*) terminator. As an exception, *LTI6b* was fused with the plant *ACTIN2* promoter (*pACT2*) for FRET-FLIM experiments (2). The BiFC YFP secreted 'SP' constructs had the LYM2 signal peptide fused to the designated YFP part.

*Transient expression in N. benthamiana*

*Agrobacterium tumefaciens* GV3101 was transformed with constructs and cultured overnight at 28 °C with appropriate antibiotics. Leaves of 4-week-old *N. benthamiana* plants were syringe-infiltrated with cultures resuspended in infiltration buffer [0.01 M 2-(N-morpholino)ethanesulfonic acid pH 5.6, 0.01 M $MgCl_2$, 0.01 M acetosyringone]. All constructs were co-infiltrated with an *Agrobacterium* strain carrying the p19 silencing suppressor, with each infiltrated at 0.5 $OD_{600nm}$ with the exception of *35S::LTI6b-mCherry* (0.2 $OD_{600nm}$). Tissue for subsequent experiments was collected or imaged two days post infiltration.

*Co-immunoprecipitation*

*N. benthamiana* leaf discs of transiently transformed tissue were frozen and homogenized. Proteins were extracted in an immunoprecipitation (IP) buffer [50 mM Tris-HCl pH 7.5, 150 mM NaCl, 1 mM EDTA, 0.5% NP40 IPEGAL® CA-630 (Sigma), 10% glycerol, protease inhibitor cocktail (Sigma) 1:100, phosphatase inhibitor (Sigma) 1:200, 1 mM $Na_2MoO_4 \times 2H_2O$, 1 mM NaF, 1.5 mM activated $Na_3VO_4$, 5 mM dithiothreitol, 1 mM PMSF]. Protein extracts were co-immunoprecipitated with RFP-Trap magnetic agarose affinity beads (Chromotek) for two hours at 4 °C with gentle agitation. Beads were washed four times with IP buffer and proteins released by heating at 70 °C in an SDS sample loading buffer containing 2-Mercaptoethanol. Proteins were separated by SDS-PAGE and transferred to an Immuno-blot® PVDF membrane.

*Extended statistical analyses*

For microprojectile bombardment assays, data were analyzed by bootstrap method (*medianBootstrap,* (3)). For callose quantifications, FRET-FLIM analysis, and rosette area measurements, a linear mixed-effects model (with independent factors specified in figure legends) was applied using the R package, *lmerTest*. For multiple images taken for a given biological replicate (*i.e.* callose quantifications and FRET-FLIM), the random effect was 'biological replicate'.

ANOVAs specified are ANOVA Satterthwaite's Method, with significant differences between factors (denoted in figure legends) determined by post hoc Tukey HSD using the R package, *emmeans.*

**Supplemental Figures and Tables**

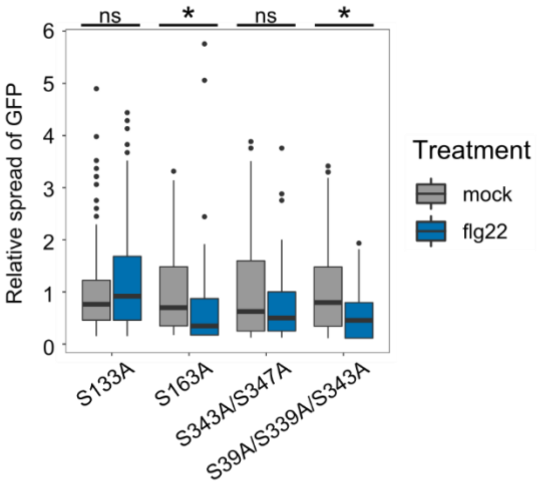


**Fig. S1 RBOHD phosphorylation is required for flg22-induced plasmodesmata closure**

Relative GFP movement to neighboring cells from microprojectile bombardment, with movement into neighboring cells reduced by flg22 in the *rbohD* mutant complemented with RBOHD$_{S163A}$ and RBOHD$_{S39A/S339A/S343A,}$ but not in the *rbohD* mutant when complemented with RBOHD$_{S133A}$ or RBOHD$_{S343A/S347A}$ (n≥ 107 bombardment sites). Asterisks indicate statistical significance compared to the mock treatment within a genotype: *$p$<0.05.

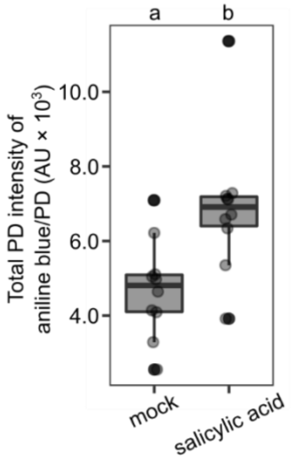


**Fig. S2 Salicylic acid induces plasmodesmata associated callose deposition within 30 minutes**

Fluorescence quantification of aniline blue stained plasmodesmata (PD)-associated callose denoted by arbitrary units (AU) using automated image analysis. 30 minutes post salicylic acid treatment increases fluorescence of callose in Col-0 *Arabidopsis*. Two z-stack image series for five biological replicates were taken per treatment. A linear mixed-effects model was used with random effect being the biological replicate. Independent factor 'Treatment' was determined as significant (ANOVA: $F = 15.2$, $df = 1$, $p<0.005$), with significant differences determined by a post hoc Tukey HSD, denoted by a and b, $p<0.005$.

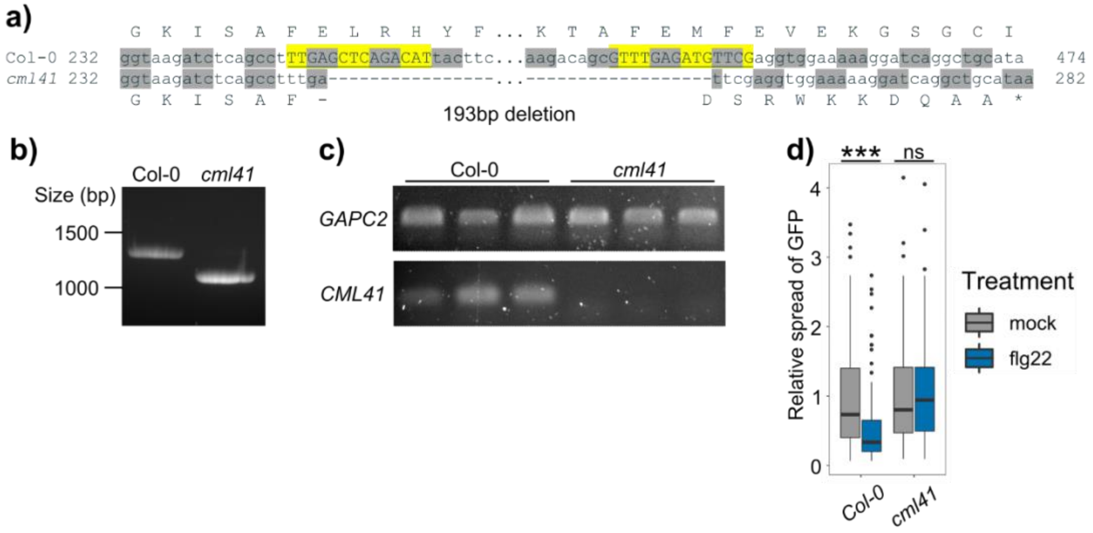


**Fig. S3 Generation of a *cml41* null mutant by CRISPR-Cas9 gene editing**

a) Target sgRNA binding sites (highlighted in yellow with sequences underlined) in the coding sequence of *CML41* with predicted gene editing introducing amino acid changes and a premature stop codon. b) A *cml41* null mutant was confirmed by detection of the gene deletion by PCR. c) No *CML41* transcript was detected in *cml41* by RT-PCR analysis with *GAPC2* expression included as a control. d) flg22 reduces GFP movement into neighboring cells in Col-0 but not in *cml41* ($n \geq 98$ bombardment sites). Data collected from four to seven biological replicates was analyzed by bootstrapping with asterisks indicating statistical significance compared to the mock treatment within a genotype: ****p*<0.001.

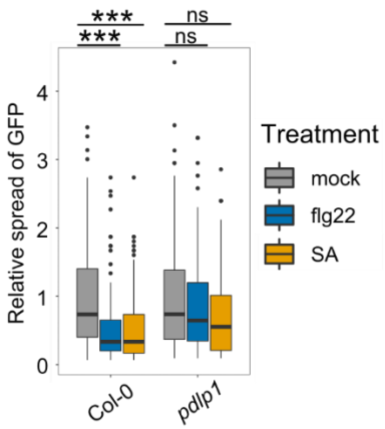


**Fig. S4 PDLP1 is required for flg22 and SA-induced plasmodesmata closure**

Relative GFP movement into neighboring cells following microprojectile bombardment. Both flg22 and SA reduce GFP movement into neighboring cells in Col-0, but not in *pdlp1* (n≥107 bombardment sites). The number of cells showing GFP has been normalized to the mean of the mock-treated data within genotypes. Asterisks indicate statistical significance compared to the mock treatment within a genotype: ***$p$<0.001.

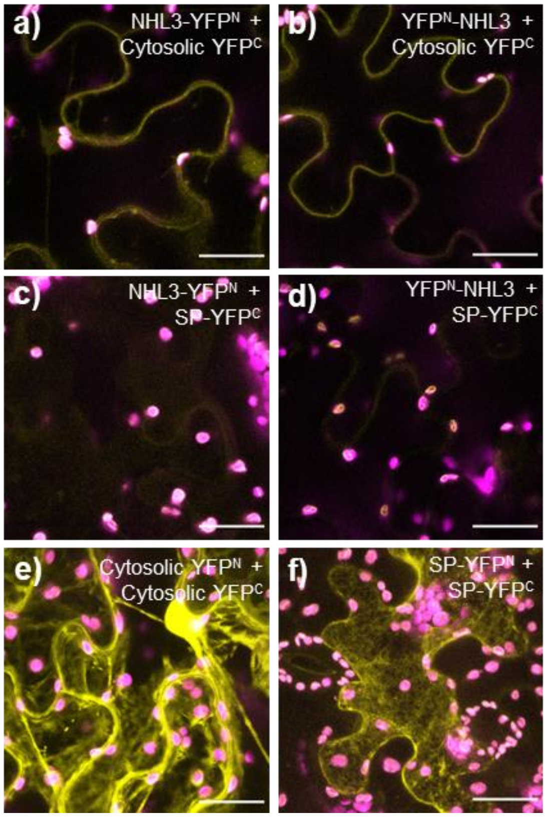


**Fig. S5 BiFC identifies that NHL3 termini are both cytosolic facing**

Maximum projections of confocal z-stacks of *N. benthamiana* epidermal cells transiently expressing split YFP constructs. The construct combinations are indicated above in each image panel, and fluorescence of split-YFP constructs is shown in yellow with chlorophyll autofluorescence in magenta. All scale bars are 25 µM.

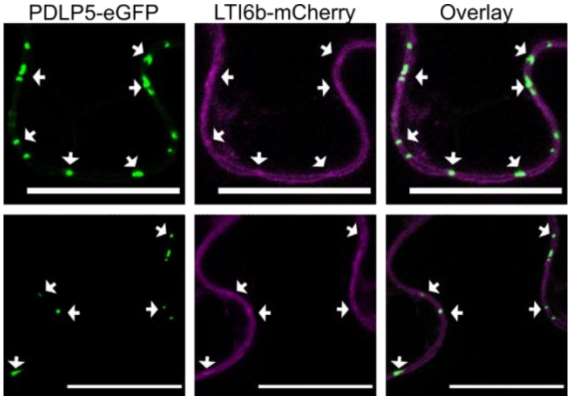


**Fig. S6 Transient expression of *pACT2::LTIb6-mCherry* identifies LTI6b-mCherry fluorescence co-incident with plasmodesmata**

Confocal micrographs of *N. benthamiana* transiently expressing *PDLP5-eGFP* and *pACT2::LTI6b-mCherry*. PDLP5-eGFP is in green and LTI6b-mCherry is in magenta. Arrows indicate co-localization at plasmodesmata. Scale bar is 25 µm.

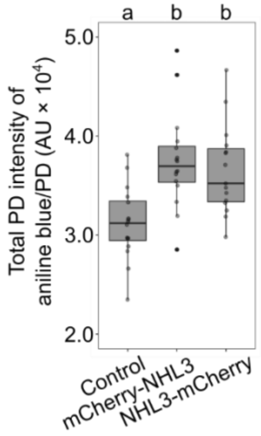


**Fig. S7 mCherry-NHL3 functions similarly to NHL3-mCherry when transiently expressed in *N. benthamiana***

Quantification of aniline blue stained plasmodesmata (PD)-associated callose denoted by arbitrary units (AU) using automated image analysis in *N. benthamiana* transiently expressing p19 only (Control), mCherry-NHL3 or NHL3-mCherry. Four z-stack image series for four biological replicates taken per construct combination. Data were analyzed by a linear mixed-effects model with random effect being the biological replicate. Independent factor ‘Gene Expressed’ was determined as significant ($F = 13.0$, $df = 2$, $p<0.0001$; ANOVA Satterthwaite’s Method), with significant differences between determined by a post hoc Tukey HSD denoted by a and b, $p<0.001$.

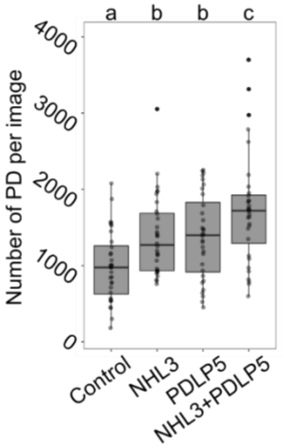


**Fig. S8 Combined expression of NHL3-mCherry and PDLP5-eGFP (NHL3+PDLP5) increases the number of plasmodesmal callose deposits detected per image**

Number of aniline blue stained plasmodesmata (PD)-associated callose using automated image analysis in *N. benthamiana* transiently expressing p19 only (Control), NHL3-mCherry (NHL3), PDLP5-eGFP (PDLP5), or NHL3-mCherry and PDLP5-eGFP (NHL3+PDLP5). Combined data from two independent experiments, with four z-stack image series from four biological replicates taken per construct combination in each experiment (for a total of 32 images per construct combination). Data were analyzed by a linear mixed-effects model with random effect being the biological replicate. Independent factor 'Gene Expressed' was determined as significant (F = 16.2, df = 3, $p$<0.0001; ANOVA Satterthwaite's Method), with significant differences determined by a post hoc Tukey HSD denoted by a, b and c, $p$<0.01.

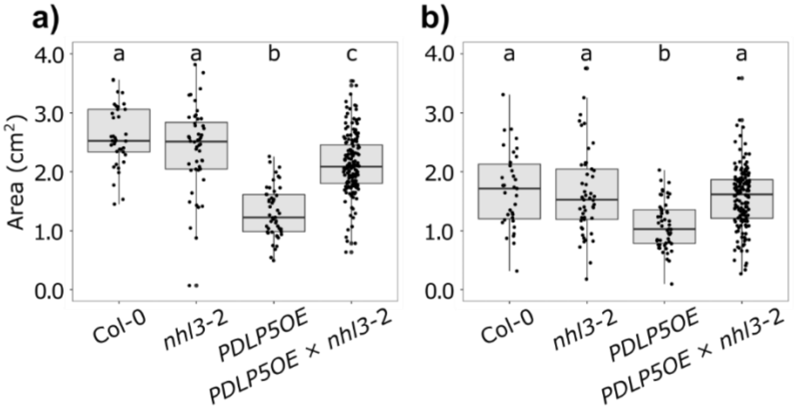


**Fig. S9 Whole rosette area of 21-day old Col-0, *nhl3*-2, *PDLP5OE* and *PDLP5OE* x *nhl3*-2**

Two independent experiments show a) partial or b) whole reversion of *PDLP5OE* dwarf phenotype. a) was analyzed by a linear mixed-effects model with random effect being the different individual plates plants were grown. Independent factor 'Genotype' was determined as significant (F = 61.8, df = 3, $p<0.0001$; ANOVA Satterthwaite's Method), with significant differences determined by a post hoc Tukey HSD denoted by a, b and c, $p<0.0001$. For Col-0, *nhl3*-2, *PDLP5OE* and *PDLP5OE* × *nhl3*-2, n = 37, 48, 49 and 160 respectively. For b), a linear mixed-effects model was used with random effects being the different individual plates plants were grown. Independent factor 'Genotype' was determined as significant (F = 24.5, df = 3, $p<0.0001$; ANOVA Satterthwaite's Method), with significant differences determined by a post hoc Tukey HSD denoted by a and b, $p<0.0001$. For Col-0, *nhl3*-2, *PDLP5OE* and *PDLP5OE* × *nhl3*-2, n = 35, 49, 51 and 158 respectively.

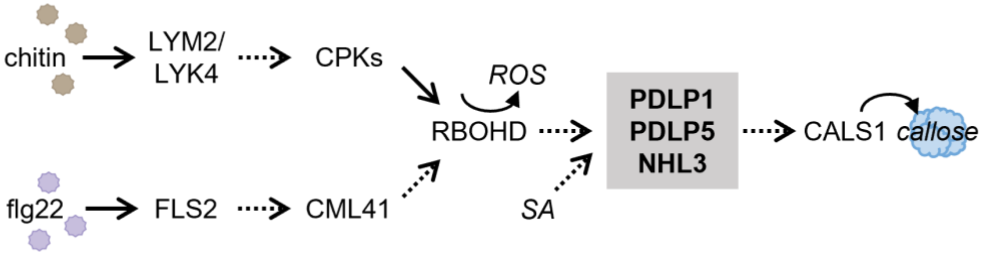


**Fig. S10 Model of elicitor induced plasmodesmal closure**

MAMP elicitors chitin and flg22 activate different receptor complexes that signal via specific pathways to RBOHD. RBOHD produces ROS that transmit plasmodesmal signals downstream, converging with SA elicited signals at a PDLP1/PDLP5/NHL3 signaling node that integrates the information and activates CALS1 to produce callose and close plasmodesmata. Established links/paths are indicated by solid arrows and yet to be understood links/paths are indicated by dashed arrows.

**Table S1 PDLP5 prey hit candidates cross-referenced with *Arabidopsis* plasmodesmal proteomes**

Split Ubiquitin Hits indicates how many protein fragments were detected in the screen. + indicates candidates found in the *Arabidopsis* Plasmodesmal Proteome (Fernandez-Calvino et al., 2011 (4)). The PD/PM ratio was taken from Brault *et al*., (2019 (5)) where its relative enrichment in the plasmodesmata fraction was compared to contaminant fractions (*i.e*., the PM, microsomal, total cell and cell wall fractions).

| Gene Identifier | Split Ubiquitin Hits | Plasmodesmal Proteome | PD/PM ratio | Name |
|---|---|---|---|---|
| **AT5G06320** | **1** | **+** | **48** | **NHL3** |
| AT1G71695 | 1 | + | | peroxidase |
| AT1G20330 | 1 | + | | SMT2 |
| AT4G23630 | 1 | + | | RTN1 |
| AT4G25810 | 1 | | 816 | XTR6 |
| AT4G35100 | 9 | + | | PIP3 |
| AT2G37170 | 3 | + | | PIP2B |
| AT1G11260 | 1 | + | | STP1 |
| AT1G53210 | 1 | + | | NCL |
| AT2G01470 | 1 | + | | STL2P |
| AT3G53420 | 1 | + | | PIP2A |
| AT3G54140 | 1 | + | | NPF8.1 |
| AT4G30190 | 1 | + | | AHA2 |
